# Female biases in islet cell gene expression, alpha cell number, and glucagon are present early in postnatal life

**DOI:** 10.64898/2026.10.27.762029

**Authors:** Carling Smith, Tiffany Z.Y. Chen, Søs Skovsø, John Ching, George Brownrigg, Janel L. Kopp, Elizabeth J. Rideout

## Abstract

Biological sex significantly affects glucose homeostasis. Sex differences in insulin sensitivity in peripheral tissues play a role in this difference; however, a growing body of evidence suggests sex differences in islet cells also contribute to differential glucose homeostasis. Studies in rodent and human models have identified sex differences in alpha– and beta-cell gene expression, in islet cell composition, metabolism, and stress responses, and in circulating levels of glucoregulatory hormones. However, most of these studies used adult humans or mice, leaving the age at which these differences emerge less clear. We performed a detailed characterization of glucose homeostasis and islet biology in young male and female mice from two mouse strains widely used in diabetes research to model both Type 1 (non-obese diabetic, NOD) and Type 2 diabetes (C57BL/6J). We uncovered male-female differences in islet cell gene expression and composition, and in circulating levels of glucagon. Importantly, these differences were reproducible across both models and were present prior to onset of diabetes-like phenotypes. Together, our data suggest sex differences in aspects of islet biology begin early in postnatal life.

## INTRODUCTION

Sex differences in glucose homeostasis are highly conserved across species (Arioglu-Inan & Kayki-Mutlu, 2023; Jo et al., 2023; Mauvais-Jarvis, 2015, 2018). For example, female individuals show lower basal circulating glucose levels and greater glucose tolerance in both rodents and humans (Arioglu-Inan & Kayki-Mutlu, 2023; Basu et al., 2006; Brownrigg et al., 2023; Jo et al., 2023; Mauvais-Jarvis, 2015, 2018). In rodent models, female individuals are also resistant to high fat diet-induced hyperglycemia (Daniels Gatward et al., 2021; Ingvorsen et al., 2017; Medrikova et al., 2012; Pettersson et al., 2012). Despite these well-known effects of biological sex on glucose homeostasis, our knowledge of the mechanisms underlying sex differences in this key trait remains incomplete due to a generalized lack of sex-specific analysis in the literature related to glucose homeostasis and diabetes (Cherian et al., 2024; Day et al., 2019).

One factor that may contribute to male-female differences in glucose homeostasis is differential tissue insulin sensitivity. In species as different as flies, mice, and humans, female individuals have greater insulin sensitivity (Biswas et al., 2025; Jo et al., 2023; Mauvais-Jarvis, 2015, 2018). While the precise molecular mechanisms are incompletely understood, greater female insulin sensitivity has been largely attributed to the insulin-sensitizing effects of estrogen (Geer & Shen, 2009; Louet et al., 2004). Beyond differential insulin sensitivity, emerging evidence shows sex differences in islet biology and islet-derived hormones may also contribute to male-female differences in glucose homeostasis. For example, adult C57BL/6J mice showed profound sex differences in islet cell gene expression, with a female bias in expression of genes associated with the unfolded protein response (UPR), endoplasmic reticulum (ER) stress, and protein synthesis (Brownrigg et al., 2023). The female bias in expression of genes within these categories was also observed in sorted murine beta cells, and in beta and alpha cells of adult human islets from donors who lived without diabetes (age 15-39) (S.-Y. Chen et al., 2026; Yong et al., 2022). Importantly, the female bias in expression of UPR-, ER stress-, and protein synthesis-related genes was associated with the greater ability of female rodent islet cells to survive and sustain glucose-stimulated insulin production during acute endoplasmic reticulum (ER) stress (Brownrigg et al., 2023).

Differential islet cell composition was recently revealed as an additional difference between male and female islets. In 3-month-old C57BL/6J mice, females showed a trend toward higher alpha cell mass compared with males, and significantly greater alpha cell mass relative to either body weight or beta cell mass compared to males (Jo et al., 2023). In 6-month-old C57BL/6J mice, female mice had lower beta cell mass and higher alpha cell area and mass compared with age-matched males (Jo et al., 2023; Visa & Berggren, 2024). Supporting these trends, islets from adult (age 15-39) human donors who lived without diabetes showed lower beta cell and higher alpha cell proportions (S.-Y. Chen et al., 2026). Additional islet parameters reported to differ between the sexes include beta cell activity in rodents (Jacobo-Piqueras et al., 2024; Visa & Berggren, 2024) and beta cell mitochondrial function in humans (S.-Y. Chen et al., 2026).

Despite this growing knowledge of male and female islets in adult individuals, it remains unclear when in development these differences are established. This knowledge is vital considering known effects of islet-derived hormones on development and diabetes pathogenesis (Cilvik et al., 2024; Jia et al., 2022; Malamitsi-Puchner et al., 2007; Rahman et al., 2021; Unger & Cherrington, 2012). We therefore compared multiple parameters of islet biology, glucose homeostasis, and circulating glucoregulatory hormones between young male and female mice in two strains of mice that are often used in diabetes research: (1) Non-obese diabetic (NOD) mice, which are frequently used to model Type 1 diabetes (T1D) mechanisms and pathogenesis; and (2) C57BL/6J, which are often used to model Type 2 diabetes (T2D) mechanisms and pathogenesis when maintained on a high fat diet (HFD) (Surwit et al., 1988; Winzell & Ahrén, 2004). Balancing our desire to compare multiple parameters between the sexes early in development, but prior to onset of T1D-like phenotypes in the NOD mice, we carried out all of our experiments in mice at approximately 4 weeks of age. While our comparison noted sex differences in several parameters that were strain-specific, female islet cells across both models showed higher expression of genes associated with UPR and ER stress, greater alpha cell number, and higher circulating glucagon levels. Sex differences in multiple islet parameters are therefore present from an early age across rodent diabetes models.

## RESULTS

### No sex difference in islet immune infiltration in young NOD mice

In pancreatic islets, immune cell infiltration may begin around 5 – 6 weeks in NOD female mice and slightly later in NOD males. Consequently, by 30 weeks of age 70-90% of female NOD mice develop T1D-like phenotypes compared to only 30-40% of NOD males (D. Chen et al., 2020). To determine a possible contribution of insulitis to sex differences in islet attributes, we quantified immune cell infiltration using CD45+ to broadly identify the presence of leukocytes. In 4-week-old NOD mice immune cell presence in the peri-islet and islet area was negligible (Fig. S1A), and we observed no sex difference in islet immune cell presence (Fig. S1A). This suggests a lack of differential islet immune cell invasion between 4-week-old male and female NOD mice.

### Female bias in expression of genes related to ER stress and UPR in young mice

Studies in adult mice and humans show profound sex differences in islet cell gene expression (Brownrigg et al., 2023; S.-Y. Chen et al., 2026; Gannon et al., 2018; Liu et al., 2021; Qadir et al., 2024; Yong et al., 2022). Using RNA sequencing, we examined gene expression in islets of 4-week-old male and female mice from NOD and C57BL/6J mice, an age at which NOD mice show no differential islet immune cell invasion and diabetes-like phenotypes are absent from both strains. After performing differential gene expression analysis (Supplemental File 5-6), we used these differentially expressed genes to perform an over-representation analysis. From genes that were upregulated in female NOD mice we uncovered 137 over-represented pathways, whereas only one pathway was over-represented in genes that were upregulated in NOD male mice (Fig. 1A; Supplementary File 5). In C57BL/6J mice, there were nine over-represented pathways from genes upregulated in C57BL/6J females and two over-represented pathways from genes upregulated in C57BL/6J males (Fig. 1B; Supplementary File 6).

**Figure 1.**
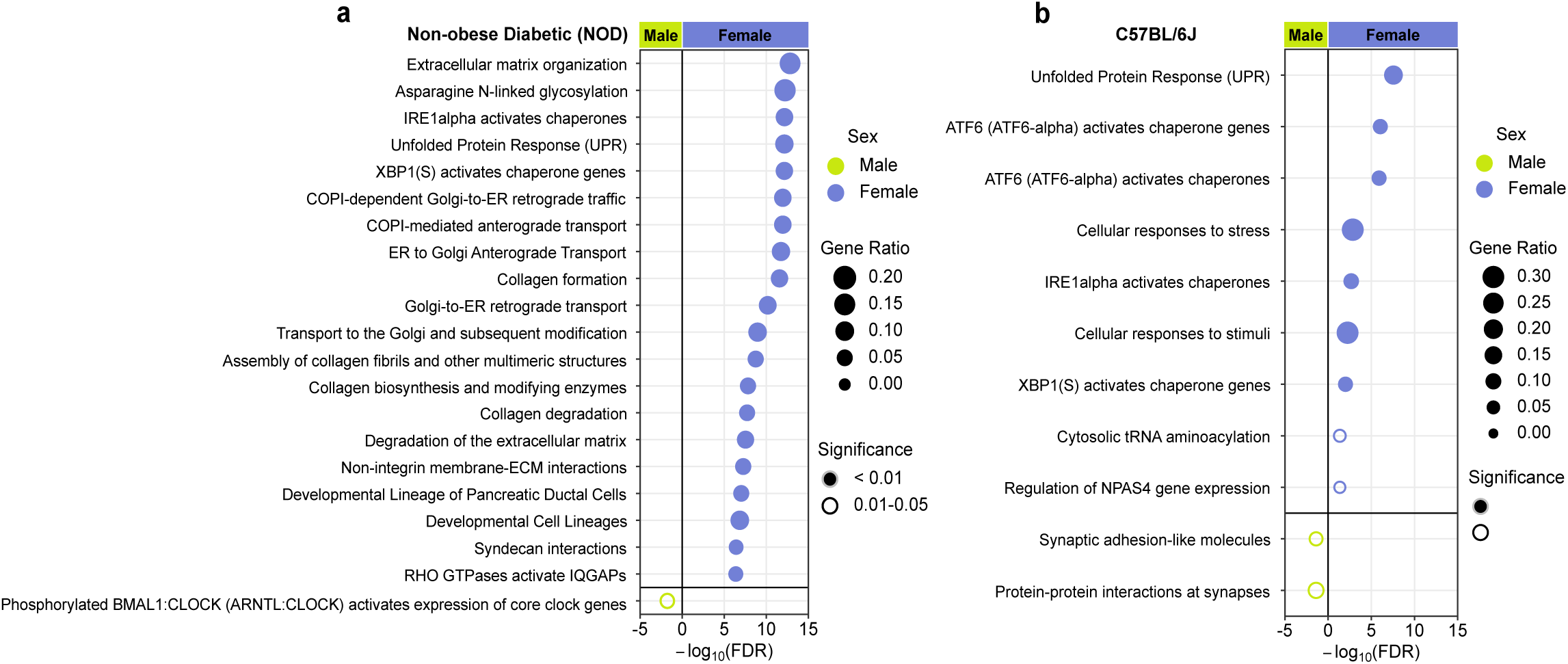
– Pathways identified as overrepresented by genes upregulated in female NOD and C57BL/6J mice. (A) Top 20/137 pathways identified as overrepresented among genes with a female bias in NOD mice, with a single pathway identified as enriched among genes with a male bias in NOD mice (*n*(f) = 10, *n*(m) = 11). (B) Nine pathways identified as overrepresented among genes with a female bias in C57BL/6J mice, with two pathways identified among genes with a male bias in C57BL/6J mice (*n*(f) = 10, (*n*(m) = 10). Full pathway details and DESEQ2 results available in Supplementary Files 1 – 6.

Overrepresented pathways with a female bias across both strains included pathways related to ER chaperones and the UPR. For example, both NOD and C57BL/6J females showed higher mRNA levels of genes associated with these pathways such as *Creld2, Pdia6, Hspa5 (Bip/Grp78), Hyou1* and *Calr*. Specifically in NOD mice, there were additional female biases in pathways related to collagen biosynthesis and modification, extracellular matrix, Golgi-ER transport, and asparagine N-linked glycosylation (Fig. 1A, Supplementary File 1). NOD males showed over-representation of a single pathway related to the circadian rhythm that included the genes *Arntl(Bmal1), Cry1, Rorc,* and *Sik1* (Fig. 1A, Supplementary File 2). Other than ER chaperones and the UPR, two pathways were uniquely over-represented in C57BL/6J females (Fig. 1A, 1B and Supplementary File 3): cytosolic tRNA aminoacylation, which supports protein synthesis, and NPAS4 gene expression, where NPAS4 may contribute to early beta cell stress responses (Sabatini et al., 2013; Yakobov et al., 2018). C57BL/6J males showed two over-represented pathways, which were related to protein interactions at synapses and synaptic adhesion-like molecules (Supplementary File 4). While not highlighted in the Reactome pathways, we noted that both NOD and C57BL/6J females had significantly higher levels of *Sst* mRNA compared to NOD and C57BL/6J males, respectively (Supplementary File 5). Together, these data show that the female bias in expression of genes related to the UPR and ER stress begins at an early age and is reproducible across rodent strains used to model T1D and T2D.

### Greater alpha cell number in young female mice

Building on these sex differences in islet gene expression, we assessed islet size and cell composition in young mice from both C57BL/6J and NOD models. We found that the average individual islet size was not significantly different between the sexes in either 4-week-old C57BL/6J or NOD mice (Fig. 2A, 2B), and the number of beta cells was equivalent between the sexes in both models at this age (Fig. 2C, 2D). In contrast, we observed a significant female bias in alpha cell number in both NOD and C57BL/6J islets (*p* = 0.022 and *p* = 0.002, respectively; Fig. 2E, 2F). Further, males contained a greater percentage of islets with no alpha cells (NOD males: 5.6%, C57BL/6J males: 13.8%) compared to female NOD and C57BL/6J mice (NOD females = 1.1%; C57BL/6J females = 4.4%). Taken together, our data indicate a female bias in alpha cell number in multiple mouse strains at a young age.

**Figure 2.**
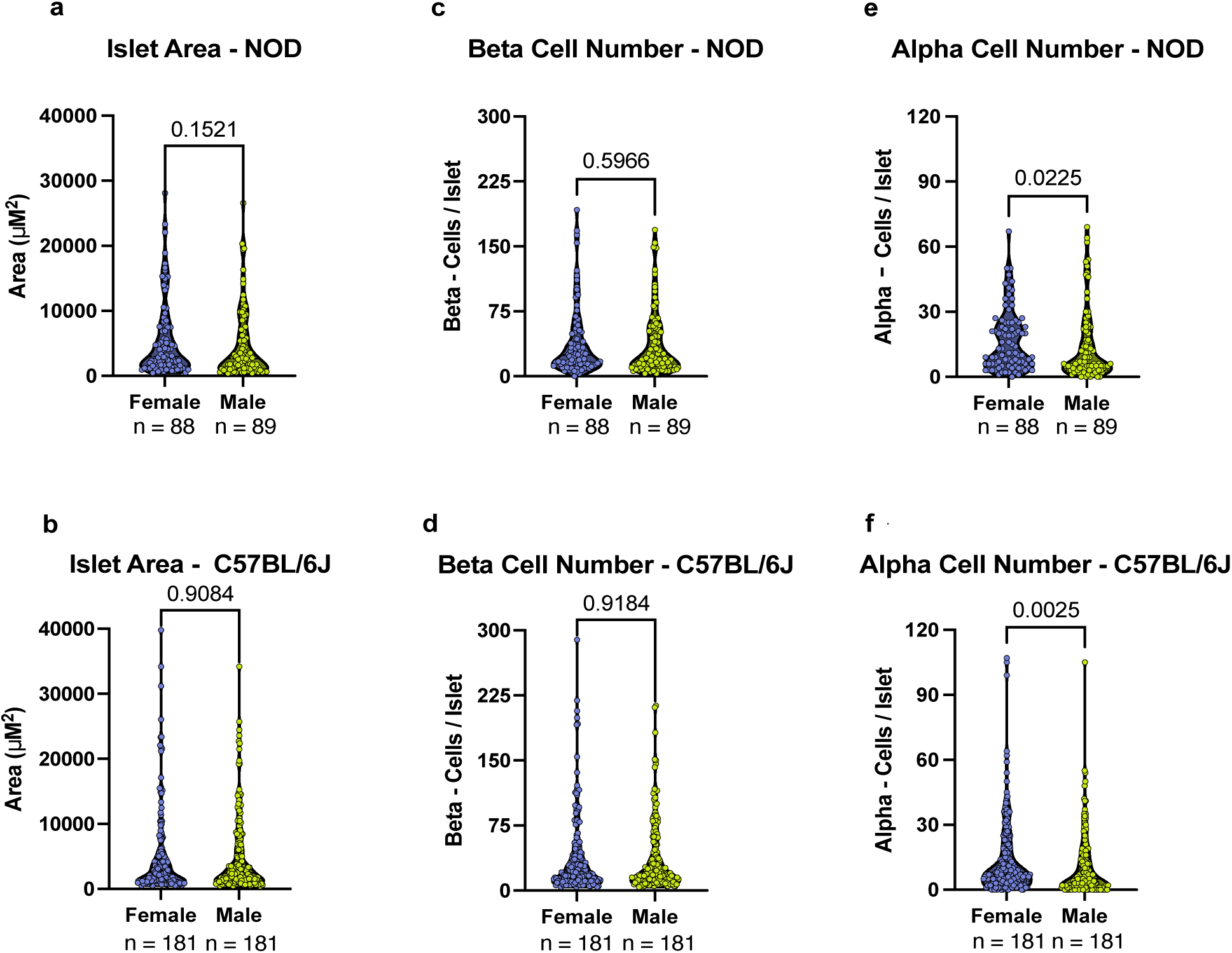
– Sex differences in pancreatic islet composition in NOD and C57BL/6J mice. Islet parameters from NOD (A-C) and C57BL/6J (D-F) mice. (A, D) Islet area did not differ between the sexes in either NOD (A; *n*(f)= 88 islets from 4 mice, *n*(m)= 89 islets from 4 mice) or C57BL/6J (D; *n*(f)= 181 islets from 5 mice, *n*(m)= 181 islets from 5 mice; Mann-Whitney test). (B, E) Beta cell number did not differ between the sexes in NOD (B; *n*(f)= 88 islets from 4 mice, *n*(m)= 89 islets from 4 mice) or C57BL/6J (E; *n*(f)= 181 islets from 5 mice, *n*(m)= 181 islets from 5 mice; Mann-Whitney test). (C, F) Alpha cell number was higher in NOD females compared to males (C; *n*(f)= 88 islets from 4 mice, *n*(m)= 89 islets from 4 mice; Mann-Whitney Test) and in C57BL/6J females (F; *n*(f)= 181 islets from 5 mice, *n*(m)= 181 islets from 5 mice; Mann-Whitney test). *n* on graphs refers to total number of islets measured. Violin plots display raw data points, with the central black line identifying the median (+/− IQR), shape displays distribution of data points.

### Higher circulating glucagon in young female mice

Based on sex differences in islet cell gene expression and composition, we compared plasma levels of insulin, proinsulin, and glucagon between males and females in 4-week-old NOD and C57BL/6J mice. In NOD mice, there was a trend toward higher plasma insulin and significantly higher plasma proinsulin levels in males compared with females (Fig. 3A, 3B); however, there was no sex difference in the proinsulin:insulin ratio (Fig. 3C). In 4-week-old C57BL/6J mice, no male-female differences were observed in either plasma insulin or proinsulin (Fig. 3D, 3E), suggesting the male biases we observed in these parameters were NOD-specific. In contrast, we found a significant female bias in plasma glucagon in 4-week-old females across both NOD and C57BL/6J models (Fig. 3G, 3H), leading to a higher glucagon:insulin ratio in both strains (Fig. S2A, S2B). Thus, the higher alpha cell number across both C57BL/6J and NOD mice aligns with higher plasma glucagon levels early in postnatal life.

**Figure 3.**
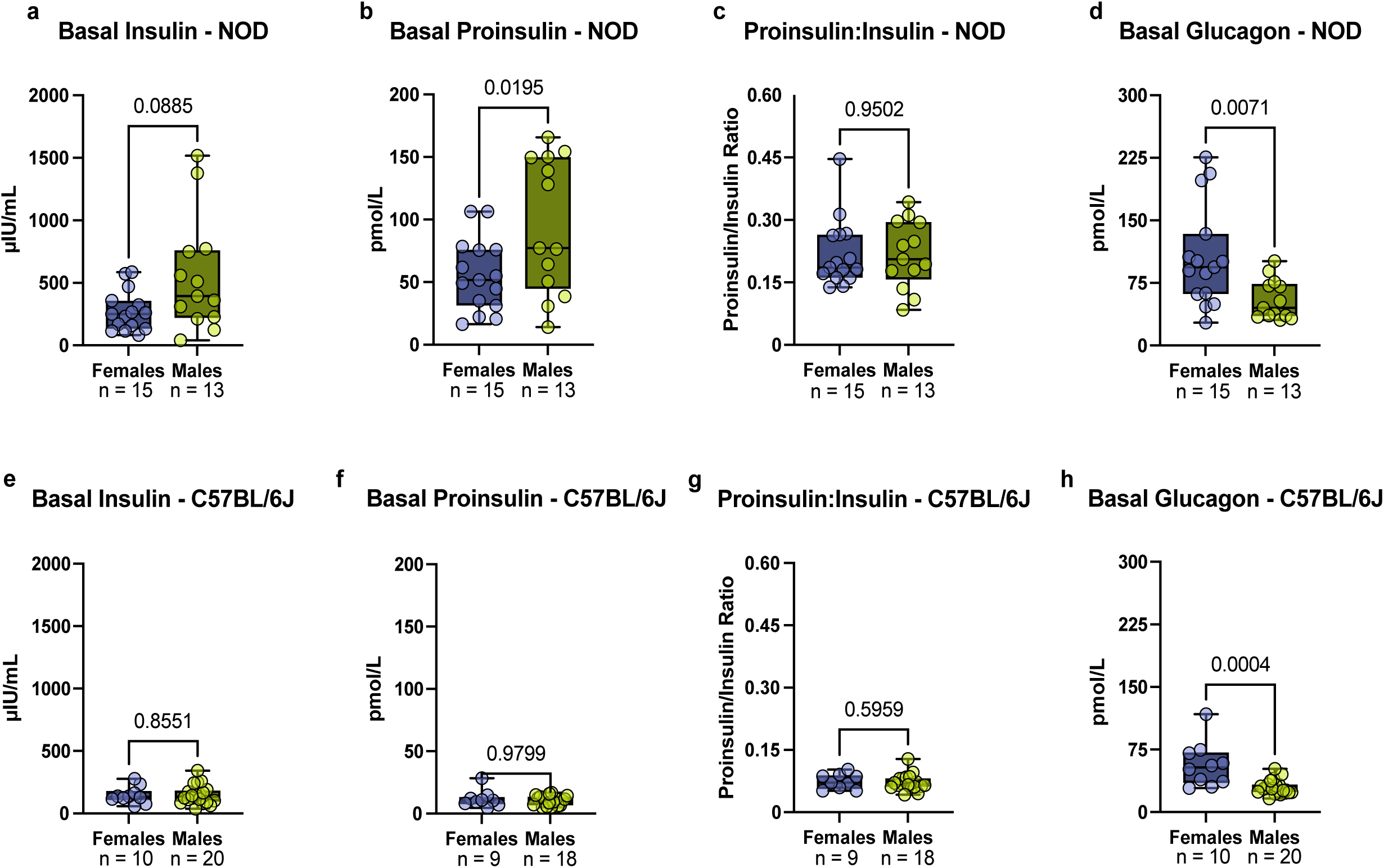
– Sex differences in glucoregulatory hormones. Glucoregulatory hormones in plasma from 4-week-old NOD (A-D) and C57BL/6J (E-G) mice. (A) A non-significant trend toward higher basal plasma insulin was found in NOD male mice (Mann-Whitney test), with significantly higher basal plasma proinsulin levels in these males compared with NOD females (B; Student’s t-test). (C) The proinsulin:insulin ratio did not differ between NOD male and female mice (Student’s t-test). (D) Basal plasma glucagon was significantly higher in female NOD mice than in males (Mann-Whitney test). Basal plasma insulin (E; Student’s t-test), proinsulin (F; Mann-Whitney test), and the proinsulin:insulin ratio (G; Mann-Whitney test) did not differ between male and female C57BL/6J mice (Student’s t-test). (H) Basal plasma glucagon was significantly higher in C57BL/6J females compared with males (Mann-Whitney test). Raw data points are displayed and median (+/− IQR) is identified as the middle bar with whiskers representing max and min values.

### Strain-specific sex differences in parameters of glucose homeostasis and regulation

To determine if sex differences in islet cell gene expression, islet composition, and production of glucoregulatory hormones influenced physiological parameters, we assessed body weight, fasting blood glucose, glucose tolerance, and insulin sensitivity in 3.3-to 4.5-week-old NOD and C57BL/6J mice (Fig. 4A-4L). In NOD mice, body weight was not significantly different between the sexes at this age (Fig. 4A) and no sex difference in fasting blood glucose was found (Fig. 4B). However, female NOD mice showed a significantly smaller increase in blood glucose at 15 minutes post-glucose injection compared to NOD males (Fig. 4C), an effect that was independent of litter, cohort or testing day, and which was apparent when blood glucose values were plotted either with or without normalization to baseline glucose (Fig. 4C, 4D and Fig. S3A). To determine if this difference was related to differential insulin sensitivity, we assessed this parameter in both sexes. There was no significant difference in insulin sensitivity between the sexes in 4-week-old NOD male and female mice (Fig. 4E, 4F).

**Figure 4.**
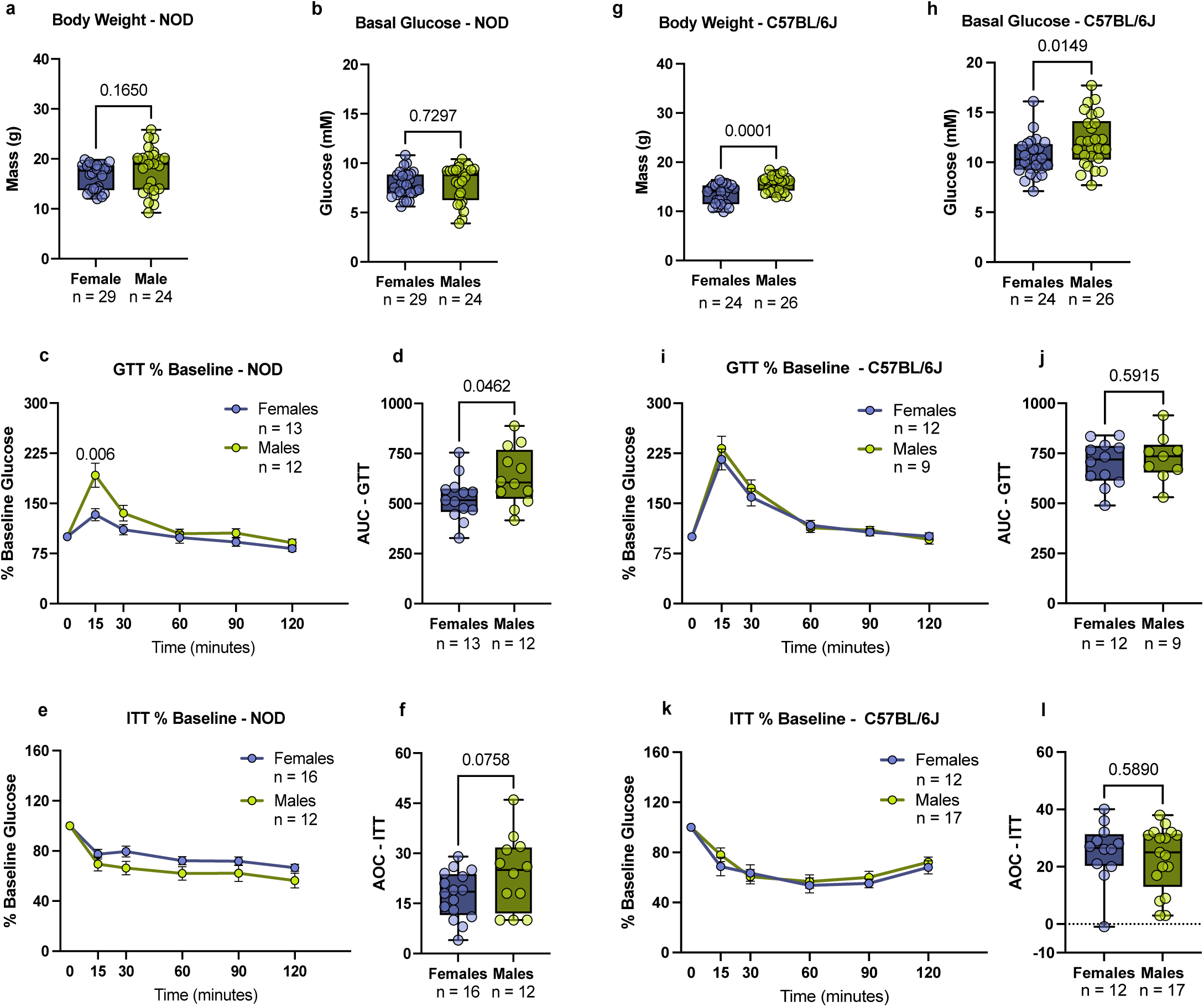
– Sex differences in percent baseline glucose and insulin tolerance, basal glucose, and body weight. Physiology measurements in female and male 4-week-old NOD (A-F) and C57BL/6J (G-L) mice. (A) Body weight (Student’s t-test) and (B) basal glucose (Student’s t-test) were not significantly different between male and female NOD mice. (C) Blood glucose levels measured after glucose injection, normalized to baseline glucose, were significantly lower in female NOD mice compared to male NOD mice at time (T) = 15 minutes (min) (Student’s t-test). (D) Total area under the curve (AUC; Student’s t-test) was significantly lower in NOD females compared to NOD males. (E) Blood glucose levels after injection of 0.75 U insulin/kg body mass, normalized to baseline glucose, were similar in NOD males and females (Students t-test). (F) Total area over the curve (AOC) for the ITT (Student’s t-test) was not significantly different between male and female NOD mice. (G) Body weight (Mann-Whitney test) and (H) basal glucose (Student’s t-test) were significantly higher in male C57BL/6J mice compared to female mice. (I) Blood glucose levels during the glucose tolerance test, normalized to baseline glucose (T = 15, 30, 60, 90, 120 min; Student’s t-test) were not significantly different. (J) Total AUC (J; Student’s t-test) was also not significantly different. (K) Neither blood glucose levels after 0.75 U insulin/kg body mass injection (T = 15, 120 min, Student’s t-test; T = 30, 60, 90 min, Mann-Whitney test) nor (L) total AOC (Student’s t-test), were significantly different between male and female C57BL/6J mice. Mean +/− SEM, error bars on GTT and ITT indicate SEM. For box plots, raw data points are displayed and median (+/− IQR) is identified as the middle bar with whiskers representing max and min values.

In 4-week-old C57BL/6J mice, body weight in males was significantly higher than in females (Fig. 4G). Fasting blood glucose was also significantly lower in female mice (Fig. 4H). While non-normalized blood glucose values suggested higher glucose tolerance in females compared with males, this difference was abolished after normalizing these values to baseline glucose (Fig. S4A and Fig. 4I). Thus, glucose tolerance in young C57BL/6J mice was equivalent between the sexes (Fig. 4I, 4J), and no sex difference in insulin sensitivity was noted at this age (Fig. 4K, 4L). Therefore, despite reproducible sex differences in islet cell gene expression, islet cell composition, and circulating glucagon levels in C57BL/6J and NOD mice, no sex differences in body weight, basal glucose, glucose tolerance, and insulin sensitivity were found in both strains.

## DISCUSSION

Our study compared islet cell gene expression and islet cell composition, glucoregulatory hormone levels, and parameters of glucose homeostasis in young male and female mice across two mouse strains frequently used to model diabetes. Even at four weeks of age, our data reveals a robust female bias in islet cell expression of genes associated with ER stress and the UPR, the number of alpha cells, and plasma glucagon levels. Due to the early age at which these differences occur, our findings have implications for the mechanisms by which sex differences in islet biology contribute to both normal physiology and diabetes pathogenesis.

For example, one key finding from our was study was a female bias in islet cell expression of ER stress– and UPR-associated genes at a very young age (*e.g*., *Bip/Grp78/Hspa5, Calr, Kdelr3, Hyou1* and *Pdia6*). CALR, HYOU1, and BiP are all forms of ER chaperone proteins (Altenhofen et al., 2023; Ikezaki et al., 2020; Yamamoto et al., 2003), and BiP is additionally known as an ER stress sensor that is essential in regulating UPR activity (Adams et al., 2019; Hartl et al., 2011; Kopp et al., 2019) (Supplementary File 1, 3, 5, 6). Given that these genes maintain a bias toward the adaptive UPR and/or are known to support islet cells in managing ER stress, female islet cells may therefore maintain this adaptive state to better manage ER stress during postnatal life (Chhabra et al., 2021; Eletto et al., 2014, 2016; Gorasia et al., 2016). Better ER stress management would be predicted to delay islet and beta cell dysfunction and the onset of diabetes in females. Consistent with this, female mice across many rodent models of chronic ER stress show delayed onset of hyperglycemia and milder islet cell defects (Austin et al., 2020; Burton et al., 2026; Daniels Gatward et al., 2021; Herbach et al., 2007; Ingvorsen et al., 2017; Pettersson et al., 2012; Winzell & Ahrén, 2004; Yoshioka et al., 1997). For example, beta cell-specific BiP loss caused greater islet cell and glucoregulatory defects in male mice than in genotype-matched females (Burton et al., 2026).

While this female bias in gene expression early in postnatal life provides a straightforward explanation for female beta cell resilience and delayed diabetes onset in mouse models of ER stress that present with T2D-like phenotypes, this explanation does not translate to the NOD model. Unlike the predominant male bias in diabetes-like phenotypes across T2D models and many human populations (Daniels Gatward et al., 2021; Jan Kakakhel et al., 2025; Kautzky-Willer et al., 2016), there is a widely-known female bias in onset of diabetes-like phenotypes in the NOD model (Pozzilli et al., 1993). Despite the fact that chronic beta cell ER stress shows a positive association with T1D pathogenesis (Engin, 2016; Engin et al., 2013; Tersey et al., 2012), as with T2D (Back & Kaufman, 2012; Engin et al., 2014; Kulkarni et al., 2023; N. Shrestha et al., 2021; Yi et al., 2020), a larger proportion of female NOD mice develop T1D-like phenotypes than males. One possible explanation for female susceptibility to T1D-like phenotypes despite better islet cell ER stress management is that a female bias in islet cell expression of other genes enhances attributes of T1D pathogenesis such as immune cell recruitment, islet infiltration, or immunogenic cell death. Genes more highly expressed in female NOD mice include genes in pathways related to MHC class II antigen presentation, cytokine signaling, interleukin signaling and antigen presentation (Supplementary File 1), and genes that have been implicated in T1D, such as *Ccl11* and *Cxcl12,* were also upregulated in female NOD mice (Cabrera et al., 2016; Chao et al., 2009; Wołoszyn-Durkiewicz et al., 2024; Zhong et al., 2024). A second possibility to explain the female bias in T1D-like phenotypes despite better ER stress management is that the main driver of this female bias in T1D lies outside of the islet. Indeed, housing conditions and the microbiome influence the incidence of T1D-like phenotypes and the sex difference in this phenotype (Markle et al., 2013; Wen et al., 2008; Yurkovetskiy et al., 2013). Future studies will therefore be needed to distinguish how similar islet cell gene expression profiles lead to a differential sex bias across two rodent strains used to model diabetes.

Another key finding from our study was that the female bias in alpha cell number and plasma glucagon was apparent early in postnatal life. Given known roles for alpha cell-derived hormones and factors (*e.g.*, GLP-1, GCG, acetylcholine and glutamate) in supporting beta cell function (Capozzi et al., 2019; Gautam et al., 2006; Gheni et al., 2014; Ho et al., 2026; Huypens et al., 2000; Y.-S. Lee & Jun, 2018; Rodriguez-Diaz et al., 2011; Svendsen et al., 2018a; Wei et al., 2023; Zhang et al., 2021), beta cell development (Habener & Stanojevic, 2012; Miller et al., 2009), regeneration of beta cell mass (Habener & Stanojevic, 2012; Wei et al., 2023) and protection against ER stress (Yusta et al., 2006), this early female bias in glucagon levels may have broad effects on glucose homeostasis and physiology. Indeed, there is extensive documentation of glucagon-mediated support for insulin secretion by stimulating beta cell glucagon and GLP-1 receptors in both mouse and human models (Capozzi et al., 2019; Ho et al., 2026; Huypens et al., 2000; Y.-S. Lee & Jun, 2018; Svendsen et al., 2018b; Wei et al., 2023; Zhang et al., 2021). Glucagon and GLP-1 receptors also enhance beta cell mass in various mouse models to maintain insulin release during HFD exposure in mice, potentially by supporting beta cell communication and coordination (Gelling et al., 2009; Suba et al., 2024; Visa & Berggren, 2024; Wei et al., 2023). Evidence also suggests that other alpha cell products such as acetylcholine and glutamate improve beta cell insulin release: the former improves beta cell glucose-sensing and the latter supports insulin granule release (Gautam et al., 2006; Gheni et al., 2014; Rodriguez-Diaz et al., 2011; Yokoi et al., 2016). Given these benefits, the loss of a female bias in alpha cell number in individuals living with T2D (S.-Y. Chen et al., 2026) suggests future studies will be needed to resolve whether glucagon contributes to sex differences in islet biology.

Finding a sex bias in islet cell biology early in postnatal life raises key questions around how this bias is established. Because these differences were present prior to onset of diabetes-like phenotypes (Fig. S1A), and in the absence of sex differences in peripheral insulin sensitivity (Fig. 4E, 4K), these factors are unlikely to drive the sex biases in islet biology. Estrogen may play a role in mediating these differences, as the female biases in gene expression and alpha cell proportion were measured around the time of the earliest known estrogen-dependent developmental events (*e.g*., vaginal opening) at approximately 4 weeks in C57BL/6J and NOD mice (Binder et al., 2023; Chester et al., 2026; Gall & Kyle, 1968; Rodriguez et al., 1997). Future studies will therefore need to determine the precise time in development when these differences arise, and comprehensively test whether estrogen, and potentially other sex-determining factors such as sex chromosome complement, contribute to this sex differences in islet biology. Importantly, both estrogen and the number of X chromosomes influence beta cell development and function as well as glucose homeostasis (Bailey & Ahmed-Sorour, 1980; A. Basu et al., 2017; Choi et al., 2005; Hall et al., 2014; Mauvais-Jarvis, 2016; Qadir et al., 2024; Wong et al., 2010).

Despite robust sex differences in islet biology and circulating glucagon levels in C57BL/6J and NOD mice, an interesting finding from our study was a lack of robust sex differences across several physiological parameters (basal glucose, glucose tolerance, body weight, insulin sensitivity). One explanation for this apparent lack of effect on glucose homeostasis relates to the limited physiological contexts we explored. For example, the impact of greater alpha cell number and higher plasma glucagon may not have been apparent in our tests because alpha cells and glucagon primarily influence these parameters after mixed-nutrient meals and during adaptation to metabolic stress induced by high-fat-diet feeding (Capozzi, Svendsen, et al., 2019; Capozzi, Wait, et al., 2019; Suba et al., 2024; Visa & Berggren, 2024). Similarly, while we saw no indication of improved glucose homeostasis in females at a young age, the cumulative effect of better ER stress management may become apparent at older ages or after metabolic challenge (Mihailidou et al., 2017; S. Shrestha et al., 2022; Yi et al., 2020). Indeed, female rodents and humans are reported to maintain more beta cell function with age and during metabolic challenges such as a high-fat diet compared to males (Bonnevie-Nielsen, 1982; Chen et al., 2026; Hrovatin et al., 2023; Li et al., 2016; Villaca et al., 2025; Visa & Berggren, 2024). Thus, despite the presence of robust sex differences in islet parameters early in postnatal life, detecting the physiological consequences of these differences requires careful consideration of the experimental or disease context in which those differences would most likely manifest.

## METHODS

### Animal care

Mice were bred in-house using breeders purchased from the Jackson Laboratory. C57BL/6J (B6J – #000664) and NOD/ShiLtJ (NOD – #001976) mice were used in all our studies. The age range used for each strain, in each experiment are as follows: for ITT NOD mice ranged from 3.3 – 4.0 weeks of age and C57BL/6J mice ranged from 3.6 – 4.1 weeks of age. For GTTs, NOD mice ranged from 4.0 – 4.5 weeks of age, and C57BL/6J mice ranged from 3.7 – 4.3 weeks of age. For basal hormone levels, NOD ages ranged from 4.0 – 4.3 weeks of age and C57BL/6J mice were all 4.1 weeks in age. RNAseq was conducted on NOD mice from 4.0 – 4.7 weeks of age and C57BL/6J mice from 4.1 – 4.9 weeks of age. Ages used for insulitis measurements in NOD mice ranged from 4.3 – 4.4 weeks of age and from 4.0 – 4.4 weeks for islet composition in NOD and C57BL/6J mice. All NOD mice were housed in the UBC Centre for Disease Modeling (CDM). C57BL/6J animals were housed in CDM for all experiments aside from islet composition where some were housed in the Modified Barrier Facility (MBF) using protocols approved by the UBC Animal Care Committee (Protocol Number: A23-0159) and in accordance with international guidelines. Mice were housed on a 12-hour light:dark cycle at 21-23°C with food and drinking water *ad libitum*, and fed a standard maintenance diet (Teklad Global Rodent Diet – Irradiated #2918 [CDM] which consisted of 18.6% energy from protein, 6.2% energy from fat, 75.2% energy from carbohydrates and Lab Diet – #5053 [MBF], 24.5% energy from protein, 13.1% energy from fat, and 62.4% energy from carbohydrates).

### Islet isolation and culture

Mouse islet isolations were performed on animals as above by ductal collagenase injection followed by filtration and hand-picking using modifications of the protocol described by Salvalaggio, as we previously published (Brownrigg et al., 2023; Salvalaggio et al., 2002). Islets recovered overnight in islet culture media (RPMI media with 11.1 mM D-glucose and supplemented with 10% v/v fetal bovine serum (FBS; Thermo, #12483020) and 1% v/v penicillin-streptomycin (P/S; Gibco, #15140-148) at 37°C with 5% CO_2_. After overnight rest, islets were washed with 1X Phosphate Buffered Saline (PBS) and resuspended in buffer RLT + beta-mercaptoethanol (BME), flash frozen on dry ice, and stored at –80°C until RNA extraction and sequencing.

### RNA sequencing

RNA was extracted using 100 islets per individual mouse, cultured as above, from both C57B6/6J (*n* = 10F, 10M) and NOD (*n* = 10F, 11M) mice. Islets were frozen in 100 *μ*L of RLT buffer (Qiagen) with BME (1%). RNA was isolated using RNeasy Mini Kit (Qiagen, #74106) according to the manufacturer’s instructions. RNA sequencing was performed at the UBC Biomedical Research Centre Sequencing Core, as previously described (Brownrigg et al, 2023). Briefly, sample quality control was performed using the Agilent 2100 Bioanalyzer System (RNA pico LabChip Kit). Samples that met quality-control standards (RIN>6.5) were prepared following the standard protocol for Illumina Stranded mRNA prep (v.2.0). Sequencing was performed on the Illumina NextSeq2000 with Paired End 59 bp x 59 bp reads. Demultiplexed read sequences were then aligned to the reference sequence (mm10) using the DRAGEN RNA app (v.4.0.4) on Basespace Sequence Hub. Raw counts were generated using htseq-count (v.2.0.5). Differential gene expression analysis was performed using the DESeq2 R package; a list of differentially-expressed genes is included as Supplementary File 5 and 6. Pathway overrepresentation analysis was performed using Reactome pathway browser (v.3.7, Database 97) using a list of differentially-expressed genes with a False Discovery Rate of < 0.05, and genes located on the sex chromosomes (X and Y) filtered out of the data set (Gillespie et al., 2022). The list of genes and their chromosome locations was acquired from NCBI (GenBank Accession No. GCA_000001635.9; Annotation: GCF_000001635.27-RS_2024_02). The list of enriched pathways is included as Supplementary File 1-4.

### In vivo analyses

Mice were fasted for 4 hours (hr) prior to glucose and insulin tolerance tests. During glucose and insulin tolerance tests, tail blood was collected for blood glucose measurements using a glucometer (One Touch Ultra 2 Glucometer, Lifescan, Canada). For all physiology tests, male and female cages were alternated to avoid effects from slightly longer fasting times at the beginning versus end of the experiment. For intraperitoneal (IP) glucose tolerance tests, the glucose dose was 2 g glucose/kg of body mass. For insulin tolerance tests, the insulin dose was 0.75 U insulin/kg body mass. For measurements of basal hormone levels, femoral blood was collected after a 4 hr fast. Blood samples were kept on ice during collection, centrifuged at 2000 x *g* for 10 minutes (min) at 4°C and stored as plasma at –80°C. Plasma samples were analyzed for insulin and glucagon using MSD plates (Cat# K152ACM) according to the manufacturer’s protocol (18151-v11-2023Dec, U-PLEX Metabolic Group 1 (mouse) Multiplex Assay, 384 well plate). Proinsulin in plasma samples was measured with ELISA kit (Mercodia, Uppsala, Sweden, Cat#10-1232-01).

### Tissue processing

The pancreatic tail and spleen were collected fresh from 4-week-old (± 2 days) male and female mice. The tissues were fixed in 4% paraformaldehyde for 24 hr at 4°C, washed 5 times with 1X PBS, and stored in 70% ethanol until processing (LEICA ASP300S). Fixed tissues were embedded using paraffin (Leica HistoCore ArcadiaH). Mouse pancreata and spleen were sectioned with a microtome.

### Immunohistochemistry

Six sections from the tail region of the pancreas, with each section separated by 70 µm, were stained with anti-CD45 immunohistochemistry as described (J. L. Kopp et al., 2012; A. Y. L. Lee et al., 2019; A. M. Y. Zhang et al., 2022). Slides were rehydrated prior to performing low pH antigen retrieval (10 mM citrate buffer at pH 6.0) in a pressure-cooker (95°C 40 min). Slides were then quenched in 4% H_2_O_2_ for 15 min. Blocking buffer was made fresh using 5% normal donkey serum in 0.1% Triton-X in 1X PBS. Slides were incubated at 4°C overnight in blocking buffer with primary rabbit monoclonal anti-CD45 antibody (1:100; Cell Signaling Cat# 70257, RRID:AB_2799780) followed by biotin-conjugated donkey anti-rabbit secondary antibody in blocking buffer (1:500; Jackson Laboratory Cat# 711-065-152, RIDD: AB_2340593). The slides were then incubated with Avidin-Biotin Complex (ABC) for 2 hr and 3,3’-diaminobenzidine (DAB) for 3 min. Lastly, Mayer’s Hematoxylin was used to stain nuclei. Slides prepared in this way were digitized using a 20x objective lens on a 3DHISTECH Panoramic MIDI slide scanner (Quorum Technologies, Guelph, Canada). Histopathological analyses were carried out using Adobe Photoshop 2024 and ImageJ (Version 1.54p). Islet borders were manually identified and outlined using Adobe Photoshop. Adobe Photoshop selection tool was used to distinguish peri-islet (180% outward from drawn islet border) from exocrine tissue. Each peri-islet area measurement was inclusive of its respective islet area. ImageJ was used to further process and quantify separated islet images: positive pixels were highlighted by adjusting the blue filter setting for every image, then the images were converted to black and white using binary function, and finally, positive area was measured. All staining, imaging, and analysis were performed without knowledge of sample parameters (*e.g*., genotype, sex).

### Immunofluorescence

Two to four tissue sections at least 70 μm apart per mouse from the islet-abundant tail region of the pancreas were immunostained for insulin and glucagon as previously described (Fu et al., 2023; A. Y. L. Lee et al., 2019). Primary antibodies included: rabbit anti-insulin (1:200; Cell Signaling Technology Cat# 3014, RRID:AB_2126503) and mouse anti-glucagon (1:100; Sigma-Aldrich Cat# G2654, RRID:AB_259852). Secondary antibodies include Cy5-conjugated AffiniPure Donkey Anti-Mouse IgG (1:200; Jackson ImmunoResearch Labs Cat# 712-175-151, RRID:AB_2619678), Alexa Fluor 488-conjugated AffiniPure Donkey Anti-Rabbit IgG (1:200; Jackson ImmunoResearch Labs Cat# 711-545-152, RRID:AB_2313584). Sections were mounted in VectaShield with 4’,6-diamidino-2-phenylindole (DAPI; SKU: VECTH1200) and Z-stacks of individual islets were captured on Leica Stellaris 5 confocal microscope at 63x magnification with oil immersion. Only intact islets with 5 or more cells were used in our analysis, where at least 20 islets were analyzed per mouse. Red, green, and blue colour channels from individual image stacks were merged in Fiji/ImageJ (1.54p) and a Z-projection was made from the stack representing 7 μm (∼1 cell thickness). Insulin– and glucagon-positive cells were manually counted using the Fiji/ImageJ cell counter add-on, and islet area was measured by manually drawing islet borders. The percentage of islets with no alpha cells was calculated by counting the number of measured islets with no visible alpha cells in the analysis image and dividing it by the total number of islets measured per that strain and sex. Counts and measurements were recorded for statistical analysis.

### Statistics

GraphPad Prism 11 (GraphPad Software, San Diego, CA, USA: GraphPad Prism) and R (v4.6.0) with RStudio were used to perform all statistical analyses and to make all graphs. R code used to generate Figure 1 was used as previously described (Brownrigg et al., 2023), changes in the code were limited to; file names, file storage locations, and graph layout/colour scheme. All data were tested for normality using the D’Agostino-Pearson test. Student’s *t*-test was the parametric test used for data with a normal distribution, whereas a Mann-Whitney test was the non-parametric test used for data that was not normally distributed. All R code is available on github: https://github.com/Rideout-Lab-UBC/R-code-for-Sex-differences-in-NOD-and-C57BL-6J-mice. Differences were considered significant if the *p*-value was less than 0.05. *: *p* < 0.05; \*\**p* < 0.01; \*\*\**p* < 0.001.

### Data Availability

RNAseq data produced in this study are publicly available (GEO: GSE338668). Raw data collected as part of this study are included in Supplemental File 7, and all statistical outputs are provided in Supplemental File 8.

## ACKNOWLEDGEMENTS

We thank members of the Rideout lab for valuable feedback and advice. We thank Xiaoke Hu from the Johnson lab for conducting pancreatic islet isolations. We thank Valkyrie Life Sciences for their help in the design and execution of physiology experiments. We thank animal care staff for supporting animal husbandry, and UBC Biomedical Research Center for performing RNA sequencing. We acknowledge that our research takes place on the traditional, ancestral, and unceded territory of the Musqueam people, a privilege for which we are grateful. This work was funded by grants to EJR from Canadian Institutes of Health Research (GS4-171365), Diabetes Canada (AWD-034225 and AWD-024145) and team grant funding from Breakthrough T1D and the Canadian Institutes of Health Research (BRET1DCA 2022).

## SUPPLEMENTAL FIGURE LEGENDS

**Figure S1 – Quantification of insulitis.** (A) CD45+ was used for generalized detection of immune cells within the peri-islet area in males and females of NOD mice. No significant difference was detected between the sexes in NOD (*n*(f) = 33 peri-islet areas, 9 mice, *n*(m) = 21 peri-islet areas, 5 mice; Mann-Whitney Test). (*n*) on the graph represents individual peri-islet areas measured. Violin plots display raw data points, with the central black line identifying the median, shape displays distribution of data points.

**Figure S2 – Glucagon:Insulin Ratio in NOD and C57BL/6J mice.** (A) Female NOD mice had significantly higher glucagon:insulin ratio compared to NOD male mice (Student’s t-test). (B) C57BL/6J females had significantly higher glucagon:insulin ratio compared to males (Student’s t-test). Raw data points are displayed and median (+/− IQR) is identified as the middle bar with whiskers representing max and min values.

**Figure S3 – Sex differences in physiology in NOD mice, non-normalized data.** (A) Female NOD mice had significantly lower blood glucose levels compared to males at T = 15, 30, 60, 90, 120 min (Student’s t-test). (B) Male and female NOD mice had similar changes in non-normalized blood glucose in response to insulin at each timepoint (Student’s t-test). Line graph data are shown as mean, error bars indicate +/− SEM. Differences were considered significant if the *p*-value was less than 0.05. *: *p* < 0.05; \*\**p* < 0.01; \*\*\**p* < 0.001.

**Figure S4 – Sex differences in physiology in C57BL/6J mice, non-normalized data.** (A-B) Glucose and insulin tolerance tests in male and female C57BL/6J mice. A) Female C57BL/6J mice had significantly lower blood glucose levels compared to males at T = 0, 15, 30, 90 min (Student’s t-test). (B) Male and female C57BL/6J mice were similarly insulin sensitive. Mean +/− SEM, error bars indicate SEM. Differences were considered significant if the *p*-value was less than 0.05. *: *p* < 0.05; \*\**p* < 0.01; \*\*\**p* < 0.001.

## REFERENCES

1. Adams, C. J., Kopp, M. C., Larburu, N., Nowak, P. R., & Ali, M. M. U. (2019). Structure and Molecular Mechanism of ER Stress Signaling by the Unfolded Protein Response Signal Activator IRE1. Frontiers in Molecular Biosciences, 6. 10.3389/fmolb.2019.00011

2. Altenhofen, D., Khuong, J. M.-A., Kuhn, T., Lebek, S., Görigk, S., Kaiser, K., Binsch, C., Griess, K., Knebel, B., Belgardt, B.-F., Cames, S., Eickelschulte, S., Stermann, T., Rasche, A., Herwig, R., Weiss, J., Vogel, H., Schürmann, A., Chadt, A., & Al-Hasani, H. (2023). E96V Mutation in the Kdelr3 Gene Is Associated with Type 2 Diabetes Susceptibility in Obese NZO Mice. International Journal of Molecular Sciences, 24(1), 845. 10.3390/ijms24010845

3. Arioglu-Inan, E., & Kayki-Mutlu, G. (2023a). Sex Differences in Glucose Homeostasis. In S. E. Tsirka & M. Acosta-Martinez (Eds.), Sex and Gender Effects in Pharmacology (pp. 219–239). Springer International Publishing. 10.1007/164_2023_664

4. Arioglu-Inan, E., & Kayki-Mutlu, G. (2023b). Sex Differences in Glucose Homeostasis. In S. E. Tsirka & M. Acosta-Martinez (Eds.), Sex and Gender Effects in Pharmacology (pp. 219–239). Springer International Publishing. 10.1007/164_2023_664

5. Austin, A. L. F., Daniels Gatward, L. F., Cnop, M., Santos, G., Andersson, D., Sharp, S., Gentry, C., Bevan, S., Jones, P. M., & King, A. J. F. (2020). The KINGS Ins2+/G32S Mouse: A Novel Model of β-Cell Endoplasmic Reticulum Stress and Human Diabetes. Diabetes, 69(12), 2667–2677. 10.2337/db20-0570

6. Back, S. H., & Kaufman, R. J. (2012). Endoplasmic Reticulum Stress and Type 2 Diabetes. Annual Review of Biochemistry, 81, 767–793. 10.1146/annurev-biochem-072909-095555

7. Bailey, C. J., & Ahmed-Sorour, H. (1980). Role of ovarian hormones in the long-term control of glucose homeostasis. Diabetologia, 19(5), 475–481. 10.1007/BF00281829

8. Basu, A., Dube, S., & Basu, R. (2017). Men Are from Mars, Women Are from Venus: Sex Differences in Insulin Action and Secretion. In F. Mauvais-Jarvis (Ed.), Sex and Gender Factors Affecting Metabolic Homeostasis, Diabetes and Obesity (pp. 53–64). Springer International Publishing. 10.1007/978-3-319-70178-3_4

9. Basu, R., Dalla Man, C., Campioni, M., Basu, A., Klee, G., Toffolo, G., Cobelli, C., & Rizza, R. A. (2006). Effects of Age and Sex on Postprandial Glucose Metabolism: Differences in Glucose Turnover, Insulin Secretion, Insulin Action, and Hepatic Insulin Extraction. Diabetes, 55(7), 2001–2014. 10.2337/db05-1692

10. Binder, A. K., Peecher, D. L., Qvigstad, A. J., Gutierrez, S. D., Magaña, J., Banks, D. B., & Korach, K. S. (2023). Differential Strain-dependent Ovarian and Metabolic Responses in a Mouse Model of PCOS. Endocrinology, 164(4), bqad024. 10.1210/endocr/bqad024

11. Biswas, P., Bako, J. A., Liston, J. B., Yu, H., Wat, L. W., Miller, C. J., Gordon, M. D., Huan, T., Stanley, M., & Rideout, E. J. (2025). Insulin/insulin-like growth factor signaling pathway promotes higher fat storage in *Drosophila* females. Cell Reports, 44(7), 115915. 10.1016/j.celrep.2025.115915

12. Bonnevie-Nielsen, V. (1982). Different effects of high glucose and high fat diet on pancreatic insulin and glucagon in female and male mice. Diabete & Metabolisme, 8(4), 271–277.

13. Brownrigg, G. P., Xia, Y. H., Chu, C. M. J., Wang, S., Chao, C., Zhang, J. A., Skovsø, S., Panzhinskiy, E., Hu, X., Johnson, J. D., & Rideout, E. J. (2023). Sex differences in islet stress responses support female β cell resilience. Molecular Metabolism, 69, 101678. 10.1016/j.molmet.2023.101678

14. Burton, J. J. N., Castro, T. A., Juray, A. A., Hassan, H. A., Diaz Mosquea, H., Sharma, R. B., & Alonso, L. C. (2026). Sex-Dependent Diabetes Impact of Acute Grp78 Deletion in β-Cells of Adult Mice. Diabetes, 75(7), 1176–1188. 10.2337/db25-0809

15. Cabrera, S. M., Henschel, A. M., & Hessner, M. J. (2016). Innate inflammation in Type 1 diabetes. Translational Research: The Journal of Laboratory and Clinical Medicine, 167(1), 214–227. 10.1016/j.trsl.2015.04.011

16. Capozzi, M. E., Svendsen, B., Encisco, S. E., Lewandowski, S. L., Martin, M. D., Lin, H., Jaffe, J. L., Coch, R. W., Haldeman, J. M., MacDonald, P. E., Merrins, M. J., D’Alessio, D. A., & Campbell, J. E. (2019). **β** Cell tone is defined by proglucagon peptides through cAMP signaling. JCI Insight, 4(5). 10.1172/jci.insight.126742

17. Capozzi, M. E., Wait, J. B., Koech, J., Gordon, A. N., Coch, R. W., Svendsen, B., Finan, B., D’Alessio, D. A., & Campbell, J. E. (2019). Glucagon lowers glycemia when **β** cells are active. JCI Insight, 4(16). 10.1172/jci.insight.129954

18. Chao, G. Y. C., Wallis, R. H., Marandi, L., & Poussier, P. (2009). Evidence for linkage of Type 1 Diabetes (T1D) resistance and upregulation of eotaxin (CCL11) in pancreatic beta cells to a chromosome 12 locus in the Biobreeding rat (49.2). The Journal of Immunology, 182(Supplement_1), 49.2. 10.4049/jimmunol.182.Supp.49.2

19. Chen, S.-Y., Cen, H. H., Chao, C. F., Pepper, A. R., Johnson, J. D., & Rideout, E. J. (2026). Biological sex affects human islet gene expression and mitochondrial function in type 2 diabetes. Endocrinology, 167(5), bqag032. 10.1210/endocr/bqag032

20. Cherian, C. M., Reeves, H. R., De Silva, D., Tsao, S., Marshall, K. E., & Rideout, E. J. (2024). Consideration of sex as a biological variable in diabetes research across twenty years. Biology of Sex Differences, 15, 19. 10.1186/s13293-024-00595-2

21. Chester, M., Wyckens, N., Airaud, E., Corre, R., Petrovic, C.-H., Pierre, A., & Guigon, C. J. (2026). Defining functional puberty in female C57BL/6J mice using endocrine, cytological and morphological markers. Scientific Reports. 10.1038/s41598-026-57314-7

22. Chhabra, N. F., Amend, A., Bastidas-Ponce, A., Sabrautzki, S., Tarquis-Medina, M., Sachs, S., Rubey, M., Lorenz-Depiereux, B., Feuchtinger, A., Bakhti, M., Lickert, H., Przemeck, G. K. H., & Hrabě de Angelis, M. (2021). A point mutation in the *Pdia6* gene results in loss of pancreatic β-cell identity causing overt diabetes. Molecular Metabolism, 54, 101334. 10.1016/j.molmet.2021.101334

23. Choi, S. B., Jang, J. S., & Park, S. (2005). Estrogen and Exercise May Enhance β-Cell Function and Mass via Insulin Receptor Substrate 2 Induction in Ovariectomized Diabetic Rats. Endocrinology, 146(11), 4786–4794. 10.1210/en.2004-1653

24. Cilvik, S. N., Boehmer, B., Wesolowski, S. R., Brown, L. D., & Rozance, P. J. (2024). Chronic late gestation fetal hyperglucagonaemia results in lower insulin secretion, pancreatic mass, islet area and beta– and α-cell proliferation. The Journal of Physiology, 602(22), 6329–6345. 10.1113/JP286974

25. Daniels Gatward, L. F., Kennard, M. R., Smith, L. I. F., & King, A. J. F. (2021). The use of mice in diabetes research: The impact of physiological characteristics, choice of model and husbandry practices. Diabetic Medicine, 38(12), e14711. 10.1111/dme.14711

26. Day, S., Wu, W., Mason, R., & Rochon, P. A. (2019). Measuring the data gap: Inclusion of sex and gender reporting in diabetes research. Research Integrity and Peer Review, 4(1), 9. 10.1186/s41073-019-0068-4

27. Eletto, D., Eletto, D., Boyle, S., & Argon, Y. (2016). PDIA6 regulates insulin secretion by selectively inhibiting the RIDD activity of IRE1. The FASEB Journal, 30(2), 653–665. 10.1096/fj.15-275883

28. Eletto, D., Eletto, D., Dersh, D., Gidalevitz, T., & Argon, Y. (2014). Protein Disulfide Isomerase A6 Controls the Decay of IRE1α Signaling via Disulfide-Dependent Association. Molecular Cell, 53(4), 562–576. 10.1016/j.molcel.2014.01.004

29. Engin, F. (2016). ER Stress and Development of Type 1 Diabetes. Journal of Investigative Medicine: The Official Publication of the American Federation for Clinical Research, 64(1), 2–6. 10.1097/JIM.0000000000000229

30. Engin, F., Nguyen, T., Yermalovich, A., & Hotamisligil, G. S. (2014). Aberrant islet unfolded protein response in type 2 diabetes. Scientific Reports, 4(1), 4054. 10.1038/srep04054

31. Engin, F., Yermalovich, A., Nguyen, T., Hummasti, S., Fu, W., Eizirik, D. L., Mathis, D., & Hotamisligil, G. S. (2013). Restoration of the Unfolded Protein Response in Pancreatic β Cells Protects Mice Against Type 1 Diabetes. Science Translational Medicine, 5(211), 211ra156-211ra156. 10.1126/scitranslmed.3006534

32. Fu, Q., Jiang, H., Qian, Y., Lv, H., Dai, H., Zhou, Y., Chen, Y., He, Y., Gao, R., Zheng, S., Liang, Y., Li, S., Xu, X., Xu, K., & Yang, T. (2023). Single-cell RNA sequencing combined with single-cell proteomics identifies the metabolic adaptation of islet cell subpopulations to high-fat diet in mice. Diabetologia, 66(4), 724–740. 10.1007/s00125-022-05849-5

33. Gall, G. A. E., & Kyle, W. H. (1968). Growth of the laboratory mouse. Theoretical and Applied Genetics, 38(7), 304–308. 10.1007/BF01297571

34. Gannon, M., Kulkarni, R. N., Tse, H. M., & Mauvais-Jarvis, F. (2018). Sex differences underlying pancreatic islet biology and its dysfunction. Molecular Metabolism, 15, 82–91. 10.1016/j.molmet.2018.05.017

35. Gautam, D., Han, S.-J., Hamdan, F. F., Jeon, J., Li, B., Li, J. H., Cui, Y., Mears, D., Lu, H., Deng, C., Heard, T., & Wess, J. (2006). A critical role for β cell M3 muscarinic acetylcholine receptors in regulating insulin release and blood glucose homeostasis in vivo. Cell Metabolism, 3(6), 449–461. 10.1016/j.cmet.2006.04.009

36. Geer, E. B., & Shen, W. (2009). Gender Differences in Insulin Resistance, Body Composition, and Energy Balance. Gender Medicine, 6(Suppl 1), 60–75. 10.1016/j.genm.2009.02.002

37. Gelling, R. W., Vuguin, P. M., Du, X. Q., Cui, L., Rømer, J., Pederson, R. A., Leiser, M., Sørensen, H., Holst, J. J., Fledelius, C., Johansen, P. B., Fleischer, N., McIntosh, C. H. S., Nishimura, E., & Charron, M. J. (2009). Pancreatic β-cell overexpression of the glucagon receptor gene results in enhanced β-cell function and mass. American Journal of Physiology-Endocrinology and Metabolism, 297(3), E695–E707. 10.1152/ajpendo.00082.2009

38. Gheni, G., Ogura, M., Iwasaki, M., Yokoi, N., Minami, K., Nakayama, Y., Harada, K., Hastoy, B., Wu, X., Takahashi, H., Kimura, K., Matsubara, T., Hoshikawa, R., Hatano, N., Sugawara, K., Shibasaki, T., Inagaki, N., Bamba, T., Mizoguchi, A., … Seino, S. (2014). Glutamate Acts as a Key Signal Linking Glucose Metabolism to Incretin/cAMP Action to Amplify Insulin Secretion. Cell Reports, 9(2), 661–673. 10.1016/j.celrep.2014.09.030

39. Gillespie, M., Jassal, B., Stephan, R., Milacic, M., Rothfels, K., Senff-Ribeiro, A., Griss, J., Sevilla, C., Matthews, L., Gong, C., Deng, C., Varusai, T., Ragueneau, E., Haider, Y., May, B., Shamovsky, V., Weiser, J., Brunson, T., Sanati, N., … D’Eustachio, P. (2022). The reactome pathway knowledgebase 2022. Nucleic Acids Research, 50(D1), D687–D692. 10.1093/nar/gkab1028

40. Gorasia, D. G., Dudek, N. L., Safavi-Hemami, H., Perez, R. A., Schittenhelm, R. B., Saunders, P. M., Wee, S., Mangum, J. E., Hubbard, M. J., & Purcell, A. W. (2016). A prominent role of PDIA6 in processing of misfolded proinsulin. Biochimica et Biophysica Acta (BBA) – Proteins and Proteomics, 1864(6), 715–723. 10.1016/j.bbapap.2016.03.002

41. Habener, J. F., & Stanojevic, V. (2012). α-cell role in β-cell generation and regeneration. Islets, 4(3), 188–198. 10.4161/isl.20500

42. Hall, E., Volkov, P., Dayeh, T., Esguerra, J. L. S., Salö, S., Eliasson, L., Rönn, T., Bacos, K., & Ling, C. (2014). Sex differences in the genome-wide DNA methylation pattern and impact on gene expression, microRNA levels and insulin secretion in human pancreatic islets. Genome Biology, 15(12), 522. 10.1186/s13059-014-0522-z

43. Hartl, F. U., Bracher, A., & Hayer-Hartl, M. (2011). Molecular chaperones in protein folding and proteostasis. Nature, 475(7356), 324–332. 10.1038/nature10317

44. Herbach, N., Rathkolb, B., Kemter, E., Pichl, L., Klaften, M., de Angelis, M. H., Halban, P. A., Wolf, E., Aigner, B., & Wanke, R. (2007). Dominant-negative effects of a novel mutated Ins2 allele causes early-onset diabetes and severe beta-cell loss in Munich Ins2C95S mutant mice. Diabetes, 56(5), 1268–1276. 10.2337/db06-0658

45. Ho, K.-H., Barmaver, S. N., Gibson, S. E., Hu, R., Yagan, M., Ahmed, H. K., Balamurugan, A. N., Jacobson, D. A., Kaverina, I., & Gu, G. (2026). Paracrine Hormonal Signals From Islet α-Cells Regulate Microtubule Dynamics in β-Cells to Promote Insulin Secretion in Mouse and Human Islets. Diabetes, 75(3), 494–505. 10.2337/db24-1025

46. Hrovatin, K., Bastidas-Ponce, A., Bakhti, M., Zappia, L., Büttner, M., Salinno, C., Sterr, M., Böttcher, A., Migliorini, A., Lickert, H., & Theis, F. J. (2023). Delineating mouse β-cell identity during lifetime and in diabetes with a single cell atlas. Nature Metabolism, 5(9), 1615–1637. 10.1038/s42255-023-00876-x

47. Huypens, P., Ling, Z., Pipeleers, D., & Schuit, F. (2000). Glucagon receptors on human islet cells contribute to glucose competence of insulin release. Diabetologia, 43(8), 1012–1019. 10.1007/s001250051484

48. Ikezaki, M., Minakata, S., Nishitsuji, K., Tabata, S., Lee Matsui, I.-S., Takatani, M., Usukura, J., Ito, Y., & Ihara, Y. (2020). Calreticulin protects insulin against reductive stress in vitro and in MIN6 cells. Biochimie, 171-172, 1–11. 10.1016/j.biochi.2020.01.011

49. Ingvorsen, C., Karp, N. A., & Lelliott, C. J. (2017). The role of sex and body weight on the metabolic effects of high-fat diet in C57BL/6N mice. Nutrition & Diabetes, 7(4), e261–e261. 10.1038/nutd.2017.6

50. Jacobo-Piqueras, N., Theiner, T., Geisler, S. M., & Tuluc, P. (2024). Molecular mechanism responsible for sex differences in electrical activity of mouse pancreatic **β** cells. JCI Insight, 9(6). 10.1172/jci.insight.171609

51. Jan Kakakhel, M. Z., Fatima, S., Ahmad, H., Ibrahim, A., Rahman, W., Haider, R., Jawad, H., Qadeer, A., Khawar, M., Ahmed, F., Ikram, J., & Kirchoff, R. W. (2025). SUN-612 Gender-Specific Prevalence and Incidence Trends in Type 1 Diabetes Mellitus: A Global Burden of Disease Analysis in Adolescents and Young Adults (Aged 15-39) from 1990 to 2021. Journal of the Endocrine Society, 9(Suppl 1), bvaf149.1261. 10.1210/jendso/bvaf149.1261

52. Jia, Y., Liu, Y., Feng, L., Sun, S., & Sun, G. (2022). Role of Glucagon and Its Receptor in the Pathogenesis of Diabetes. Frontiers in Endocrinology, 13. 10.3389/fendo.2022.928016

53. Jo, S., Beetch, M., Gustafson, E., Wong, A., Oribamise, E., Chung, G., Vadrevu, S., Satin, L. S., Bernal-Mizrachi, E., & Alejandro, E. U. (2023). Sex Differences in Pancreatic β-Cell Physiology and Glucose Homeostasis in C57BL/6J Mice. Journal of the Endocrine Society, 7(9), bvad099. 10.1210/jendso/bvad099

54. Kautzky-Willer, A., Leutner, M., & Harreiter, J. (2023). Sex differences in type 2 diabetes. Diabetologia, 66(6), 986–1002. 10.1007/s00125-023-05891-x

55. Kopp, J. L., von Figura, G., Mayes, E., Liu, F.-F., Dubois, C. L., Morris, J. P., Pan, F. C., Akiyama, H., Wright, C. V. E., Jensen, K., Hebrok, M., & Sander, M. (2012). Identification of Sox9-dependent acinar-to-ductal reprogramming as the principal mechanism for initiation of pancreatic ductal adenocarcinoma. Cancer Cell, 22(6), 737–750. 10.1016/j.ccr.2012.10.025

56. Kopp, M. C., Larburu, N., Durairaj, V., Adams, C. J., & Ali, M. M. U. (2019). UPR proteins IRE1 and PERK switch BiP from chaperone to ER stress sensor. Nature Structural & Molecular Biology, 26(11), 1053–1062. 10.1038/s41594-019-0324-9

57. Kulkarni, A., Muralidharan, C., May, S. C., Tersey, S. A., & Mirmira, R. G. (2023). Inside the β Cell: Molecular Stress Response Pathways in Diabetes Pathogenesis. Endocrinology, 164(1), bqac184. 10.1210/endocr/bqac184

58. Lee, A. Y. L., Dubois, C. L., Sarai, K., Zarei, S., Schaeffer, D. F., Sander, M., & Kopp, J. L. (2019). Cell of origin affects tumor development and phenotype in pancreatic ductal adenocarcinoma. Gut, 68(3), 487–498. 10.1136/gutjnl-2017-314426

59. Lee, Y.-S., & Jun, H.-S. (2018). Glucagon-Like Peptide-1 Receptor Agonist and Glucagon Increase Glucose-Stimulated Insulin Secretion in Beta Cells via Distinct Adenylyl Cyclases. International Journal of Medical Sciences, 15(6), 603–609. 10.7150/ijms.24492

60. Li, L., El-Kholy, W., Rhodes, C. J., & Brubaker, P. L. (2005). Glucagon-like peptide-1 protects beta cells from cytokine-induced apoptosis and necrosis: Role of protein kinase B. Diabetologia, 48(7), 1339–1349. 10.1007/s00125-005-1787-2

61. Li, T., Jiao, W., Li, W., & Li, H. (2016). Sex effect on insulin secretion and mitochondrial function in pancreatic beta cells of elderly Wistar rats. Endocrine Research, 41(3), 167–179. 10.3109/07435800.2015.1124437

62. Liu, G., Li, Y., Zhang, T., Li, M., Li, S., He, Q., Liu, S., Xu, M., Xiao, T., Shao, Z., Shi, W., & Li, W. (2021). Single-Cell RNA Sequencing Reveals Sexually Dimorphic Transcriptome and Type 2 Diabetes Genes in Mouse Islet β Cells. *Genomics*, Proteomics & Bioinformatics, 19(3), 408–422. 10.1016/j.gpb.2021.07.004

63. Louet, J.-F., LeMay, C., & Mauvais-Jarvis, F. (2004). Antidiabetic actions of estrogen: Insight from human and genetic mouse models. Current Atherosclerosis Reports, 6(3), 180–185. 10.1007/s11883-004-0030-9

64. Malamitsi-Puchner, A., Briana, D. D., Gourgiotis, D., Boutsikou, M., Puchner, K.-P., Baka, S., Marmarinos, A., & Hassiakos, D. (2007). Insulin-Like Growth Factor (IGF)-I and Insulin in Normal and Growth-Restricted Mother/Infant Pairs. Mediators of Inflammation, 2007, 42646. 10.1155/2007/42646

65. Markle, J. G. M., Frank, D. N., Mortin-Toth, S., Robertson, C. E., Feazel, L. M., Rolle-Kampczyk, U., von Bergen, M., McCoy, K. D., Macpherson, A. J., & Danska, J. S. (2013). Sex differences in the gut microbiome drive hormone-dependent regulation of autoimmunity. Science, 339(6123), 1084–1088. 10.1126/science.1233521

66. Mauvais-Jarvis, F. (2015). Sex differences in metabolic homeostasis, diabetes, and obesity. Biology of Sex Differences, 6, 14. 10.1186/s13293-015-0033-y

67. Mauvais-Jarvis, F. (2016). Role of sex steroids in β-cell function, growth and survival. Trends in Endocrinology and Metabolism: TEM, 27(12), 844–855. 10.1016/j.tem.2016.08.008

68. Mauvais-Jarvis, F. (2018). Gender differences in glucose homeostasis and diabetes. Physiology & Behavior, The Proceedings of the American University Symposium on Sex Differences: From Neuroscience to the Clinic and Beyond, 187, 20–23. 10.1016/j.physbeh.2017.08.016

69. Medrikova, D., Jilkova, Z. M., Bardova, K., Janovska, P., Rossmeisl, M., & Kopecky, J. (2012). Sex differences during the course of diet-induced obesity in mice: Adipose tissue expandability and glycemic control. International Journal of Obesity, 36(2), 262–272. 10.1038/ijo.2011.87

70. Mihailidou, C., Chatzistamou, I., Papavassiliou, A. G., & Kiaris, H. (2017). Modulation of Pancreatic Islets’ Function and Survival During Aging Involves the Differential Regulation of Endoplasmic Reticulum Stress by p21 and CHOP. Antioxidants & Redox Signaling, 27(4), 185–200. 10.1089/ars.2016.6671

71. Miller, K., Kim, A., Kilimnik, G., Jo, J., Moka, U., Periwal, V., & Hara, M. (2009). Islet Formation during the Neonatal Development in Mice. PLoS ONE, 4(11), e7739. 10.1371/journal.pone.0007739

72. Mullen, Y. (2017). Development of the Nonobese Diabetic Mouse and Contribution of Animal Models for Understanding Type 1 Diabetes. Pancreas, 46(4), 455–466. 10.1097/MPA.0000000000000828

73. Pettersson, U. S., Waldén, T. B., Carlsson, P.-O., Jansson, L., & Phillipson, M. (2012). Female Mice are Protected against High-Fat Diet Induced Metabolic Syndrome and Increase the Regulatory T Cell Population in Adipose Tissue. PLOS ONE, 7(9), e46057. 10.1371/journal.pone.0046057

74. Pozzilli, P., Signore, A., Williams, A. J. K., & Beales, P. E. (1993). NOD mouse colonies around the world-recent facts and figures. Immunology Today, 14(5), 193–196. 10.1016/0167-5699(93)90160-M

75. Qadir, M. M. F., Elgamal, R. M., Song, K., Kudtarkar, P., Sakamuri, S. S. V. P., Katakam, P. V., El-Dahr, S. S., Kolls, J. K., Gaulton, K. J., & Mauvais-Jarvis, F. (2024). Sex-specific regulatory architecture of pancreatic islets from subjects with and without type 2 diabetes. The EMBO Journal, 43(24), 6364–6382. 10.1038/s44318-024-00313-z

76. Rahman, M. S., Hossain, K. S., Das, S., Kundu, S., Adegoke, E. O., Rahman, Md. A., Hannan, Md. A., Uddin, M. J., & Pang, M.-G. (2021). Role of Insulin in Health and Disease: An Update. International Journal of Molecular Sciences, 22(12), 6403. 10.3390/ijms22126403

77. Reddy, S., Liu, W., Thompson, J. M. D., Bibby, N. J., & Elliott, R. B. (1992). First phase insulin release in the non-obese diabetic mouse: Correlation with insulitis, beta cell number and autoantibodies. Diabetes Research and Clinical Practice, 17(1), 17–25. 10.1016/0168-8227(92)90039-T

78. Rodriguez, I., Araki, K., Khatib, K., Martinou, J.-C., & Vassalli, P. (1997). Mouse Vaginal Opening Is an Apoptosis-Dependent Process Which Can Be Prevented by the Overexpression of Bcl2. Developmental Biology, 184(1), 115–121. 10.1006/dbio.1997.8522

79. Rodriguez-Diaz, R., Dando, R., Jacques-Silva, M. C., Fachado, A., Molina, J., Abdulreda, M. H., Ricordi, C., Roper, S. D., Berggren, P.-O., & Caicedo, A. (2011). Alpha cells secrete acetylcholine as a non-neuronal paracrine signal priming beta cell function in humans. Nature Medicine, 17(7), 888–892. 10.1038/nm.2371

80. Sabatini, P. V., Krentz, N. A. J., Zarrouki, B., Westwell-Roper, C. Y., Nian, C., Uy, R. A., Shapiro, A. M. J., Poitout, V., & Lynn, F. C. (2013). Npas4 Is a Novel Activity–Regulated Cytoprotective Factor in Pancreatic β-Cells. Diabetes, 62(8), 2808–2820. 10.2337/db12-1527

81. Salvalaggio, P. R. O., Deng, S., Ariyan, C. E., Millet, I., Zawalich, W. S., Basadonna, G. P., & Rothstein, D. M. (2002). Islet filtration: A simple and rapid new purification procedure that avoids ficoll and improves islet mass and function. Transplantation, 74(6), 877. https://journals.lww.com/transplantjournal/pages/articleviewer.aspx?year=2002&issue=09270&article=00023&type=Fulltext

82. Shrestha, N., De Franco, E., Arvan, P., & Cnop, M. (2021). Pathological β-Cell Endoplasmic Reticulum Stress in Type 2 Diabetes: Current Evidence. Frontiers in Endocrinology, 12. 10.3389/fendo.2021.650158

83. Shrestha, S., Erikson, G., Lyon, J., Spigelman, A. F., Bautista, A., Manning Fox, J. E., dos Santos, C., Shokhirev, M., Cartailler, J.-P., Hetzer, M. W., MacDonald, P. E., & Arrojo e Drigo, R. (2022). Aging compromises human islet beta cell function and identity by decreasing transcription factor activity and inducing ER stress. Science Advances, 8(40), eabo3932. 10.1126/sciadv.abo3932

84. Signore, A., Procaccini, E., Toscano, A. M., Ferretti, E., Pozzilli, P., Williams, A. J. K., Beales, P. E., & Cugini, P. (1994). Histological study of pancreatic beta-cell loss in relation to the insulitis process in the non-obese diabetic mouse. Histochemistry, 101(4), 263–269. 10.1007/BF00315913

85. Suba, K., Patel, Y., Martin-Alonso, A., Hansen, B., Xu, X., Roberts, A., Norton, M., Chung, P., Shrewsbury, J., Kwok, R., Kalogianni, V., Chen, S., Liu, X., Kalyviotis, K., Rutter, G. A., Jones, B., Minnion, J., Owen, B. M., Pantazis, P., … Salem, V. (2024). Intra-islet glucagon signalling regulates beta-cell connectivity, first-phase insulin secretion and glucose homoeostasis. Molecular Metabolism, 85, 101947. 10.1016/j.molmet.2024.101947

86. Svendsen, B., Larsen, O., Gabe, M. B. N., Christiansen, C. B., Rosenkilde, M. M., Drucker, D. J., & Holst, J. J. (2018). Insulin Secretion Depends on Intra-islet Glucagon Signaling. Cell Reports, 25(5), 1127–1134.e2. 10.1016/j.celrep.2018.10.018

87. Tersey, S. A., Nishiki, Y., Templin, A. T., Cabrera, S. M., Stull, N. D., Colvin, S. C., Evans-Molina, C., Rickus, J. L., Maier, B., & Mirmira, R. G. (2012). Islet β-Cell Endoplasmic Reticulum Stress Precedes the Onset of Type 1 Diabetes in the Nonobese Diabetic Mouse Model. Diabetes, 61(4), 818–827. 10.2337/db11-1293

88. Unger, R. H., & Cherrington, A. D. (2012). Glucagonocentric restructuring of diabetes: A pathophysiologic and therapeutic makeover. The Journal of Clinical Investigation, 122(1), 4–12. 10.1172/JCI60016

89. Villaca, C. B. P., Oliveira, V. R., Santos, G. J., & Ortis, F. (2025). The effect of HNF4α knockout in beta cells is age and sex dependent. Islets, 17(1), 2552549. 10.1080/19382014.2025.2552549

90. Visa, M., & Berggren, P.-O. (2024). Sex-dependent intra-islet structural rearrangements affecting alpha-to-beta cell interactions lead to adaptive enhancements of Ca2+ dynamics in prediabetic beta cells. Diabetologia, 67(8), 1663–1682. 10.1007/s00125-024-06173-w

91. Wei, T., Cui, X., Jiang, Y., Wang, K., Wang, D., Li, F., Lin, X., Gu, L., Yang, K., Yang, J., Hong, T., & Wei, R. (2023). Glucagon Acting at the GLP-1 Receptor Contributes to β-Cell Regeneration Induced by Glucagon Receptor Antagonism in Diabetic Mice. Diabetes, 72(5), 599–610. 10.2337/db22-0784

92. Wen, L., Ley, R. E., Volchkov, P. Y., Stranges, P. B., Avanesyan, L., Stonebraker, A. C., Hu, C., Wong, F. S., Szot, G. L., Bluestone, J. A., Gordon, J. I., & Chervonsky, A. V. (2008). Innate immunity and intestinal microbiota in the development of Type 1 diabetes. Nature, 455(7216), 1109–1113. 10.1038/nature07336

93. Winzell, M. S., & Ahrén, B. (2004). The high-fat diet-fed mouse: A model for studying mechanisms and treatment of impaired glucose tolerance and type 2 diabetes. Diabetes, 53 *Suppl 3*, S215–219. 10.2337/diabetes.53.suppl_3.s215

94. Wołoszyn-Durkiewicz, A., Iwaszkiewicz-Grześ, D., Świętoń, D., Kujawa, M. J., Jankowska, A., Durawa, A., Glasner, P., Trzonkowski, P., Glasner, L., Szurowska, E., & Myśliwiec, M. (2024). The Complex Network of Cytokines and Chemokines in Pediatric Patients with Long-Standing Type 1 Diabetes. International Journal of Molecular Sciences, 25(3), 1565. 10.3390/ijms25031565

95. Wong, W. P. S., Tiano, J. P., Liu, S., Hewitt, S. C., Le May, C., Dalle, S., Katzenellenbogen, J. A., Katzenellenbogen, B. S., Korach, K. S., & Mauvais-Jarvis, F. (2010). Extranuclear estrogen receptor-α stimulates NeuroD1 binding to the insulin promoter and favors insulin synthesis. Proceedings of the National Academy of Sciences, 107(29), 13057–13062. 10.1073/pnas.0914501107

96. Yakobov, N., Debard, S., Fischer, F., Senger, B., & Becker, H. D. (2018). Cytosolic aminoacyl-tRNA synthetases: Unanticipated relocations for unexpected functions. *Biochimica et Biophysica Acta (BBA) – Gene Regulatory Mechanisms*, Regulation of tRNA Synthesis and Modification in Physiological Conditions and Disease, 1861(4), 387–400. 10.1016/j.bbagrm.2017.11.004

97. Yamamoto, K., Hamada, H., Shinkai, H., Kohno, Y., Koseki, H., & Aoe, T. (2003). The KDEL Receptor Modulates the Endoplasmic Reticulum Stress Response through Mitogen-activated Protein Kinase Signaling Cascades*. Journal of Biological Chemistry, 278(36), 34525–34532. 10.1074/jbc.M304188200

98. Yi, X., Cai, X., Wang, S., & Xiao, Y. (2020). Mechanisms of impaired pancreatic β-cell function in high-fat diet-induced obese mice: The role of endoplasmic reticulum stress. Molecular Medicine Reports, 21(5), 2041–2050. 10.3892/mmr.2020.11013

99. Yokoi, N., Gheni, G., Takahashi, H., & Seino, S. (2016). β-Cell glutamate signaling: Its role in incretin-induced insulin secretion. Journal of Diabetes Investigation, 7(Suppl 1), 38–43. 10.1111/jdi.12468

100. Yong, H. J., Toledo, M. P., Nowakowski, R. S., & Wang, Y. J. (2022). Sex Differences in the Molecular Programs of Pancreatic Cells Contribute to the Differential Risks of Type 2 Diabetes. Endocrinology, 163(11), bqac156. 10.1210/endocr/bqac156

101. Yoshioka, M., Kayo, T., Ikeda, T., & Koizumi, A. (1997). A novel locus, Mody4, distal to D7Mit189 on chromosome 7 determines early-onset NIDDM in nonobese C57BL/6 (Akita) mutant mice. Diabetes, 46(5), 887–894. 10.2337/diab.46.5.887

102. Yurkovetskiy, L., Burrows, M., Khan, A. A., Graham, L., Volchkov, P., Becker, L., Antonopoulos, D., Umesaki, Y., & Chervonsky, A. V. (2013). Gender bias in autoimmunity is influenced by microbiota. Immunity, 39(2), 10.1016/j.immuni.2013.08.013. https://doi.org/10.1016/j.immuni.2013.08.013

103. Yusta, B., Baggio, L. L., Estall, J. L., Koehler, J. A., Holland, D. P., Li, H., Pipeleers, D., Ling, Z., & Drucker, D. J. (2006). GLP-1 receptor activation improves β cell function and survival following induction of endoplasmic reticulum stress. Cell Metabolism, 4(5), 391–406. 10.1016/j.cmet.2006.10.001

104. Zhang, A. M. Y., Chu, K. H., Daly, B. F., Ruiter, T., Dou, Y., Yang, J. C. C., de Winter, T. J. J., Chhuor, J., Wang, S., Flibotte, S., Zhao, Y. B., Hu, X., Li, H., Rideout, E. J., Schaeffer, D. F., Johnson, J. D., & Kopp, J. L. (2022). Effects of hyperinsulinemia on pancreatic cancer development and the immune microenvironment revealed through single-cell transcriptomics. Cancer & Metabolism, 10, 5. 10.1186/s40170-022-00282-z

105. Zhang, Y., Han, C., Zhu, W., Yang, G., Peng, X., Mehta, S., Zhang, J., Chen, L., & Liu, Y. (2021). Glucagon Potentiates Insulin Secretion Via β-Cell GCGR at Physiological Concentrations of Glucose. Cells, 10(9), 2495. 10.3390/cells10092495

106. Zhong, T., Li, X., Lei, K., Tang, R., Zhou, Z., Zhao, B., & Li, X. (2024). CXCL12-CXCR4 mediates CD57+ CD8+ T cell responses in the progression of type 1 diabetes. Journal of Autoimmunity, 143, 103171. 10.1016/j.jaut.2024.103171

