## Supplementary figures and images for "Female biases in islet cell gene expression, alpha cell number, and glucagon are present early in postnatal life"

### Supplemental Figure 1

**Supplementary Figure 1.**

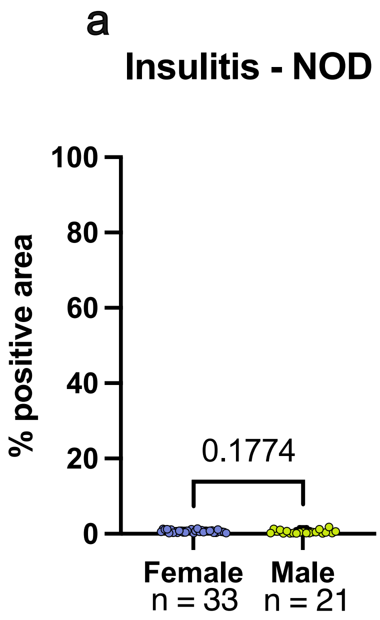

### Supplemental Figure 2

**Supplementary Figure 2.**

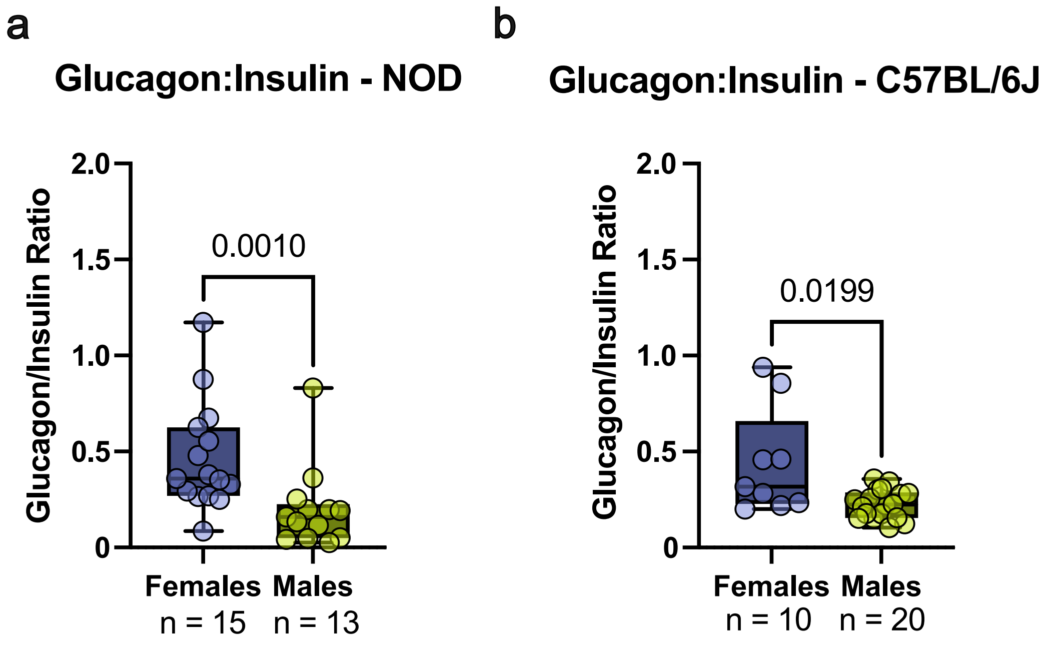

### Supplemental Figure 3

**Supplementary Figure 3.**

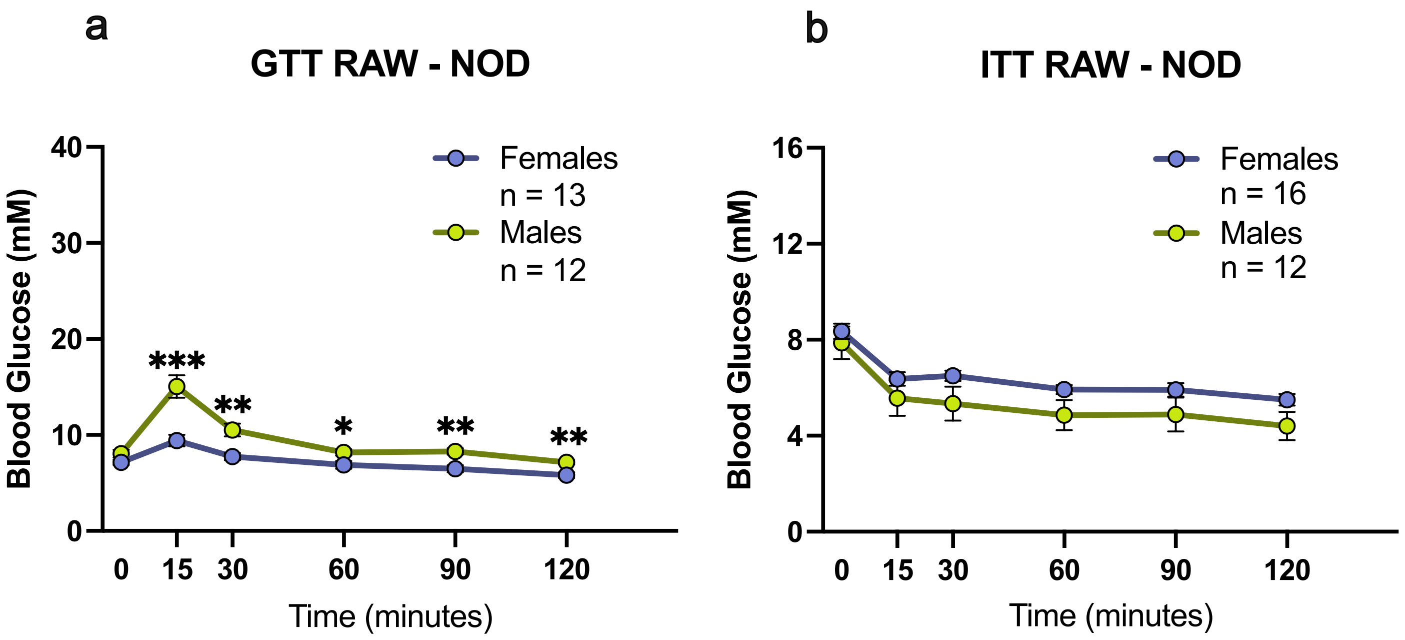

### Supplemental Figure 4

Supplementary Figure 4.

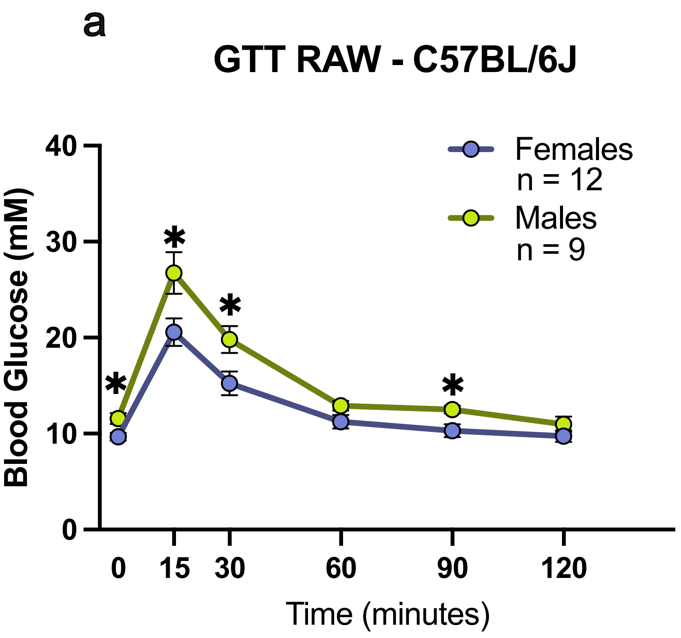
